# *Latrophilin-3* conditional knockout in tyrosine hydroxylase neurons (*Lphn3-Th-Cre*) Compared with *Lphn3* Global KO rats: Role of *Lphn3* in Tyrosine Hydroxylase Neurons on the Cognitive and Behavioral Effects of this ADHD Susceptibility Gene

**DOI:** 10.1101/2024.12.27.630427

**Authors:** Samantha L. Regan, Chiho Sugimoto, Aliyah N. Lingo, Erin A. Tepe, Michael T. Williams, Charles V. Vorhees

## Abstract

Latrophilin-3 (LPHN3) is a brain specific adhesion G-protein coupled receptor associated with elevated risk of attention deficit hyperactivity disorder (ADHD). We developed a global *Lphn3* knock-out (gKO) rat using CRISPR/Cas9 to delete exon-3. Here we report the development of a floxed *Lphn3* rat crossed with tyrosine hydroxylase (*Th-Cre*) rats to create a conditional *Lphn3* KO rat specific for catecholaminergic- positive cells. The gKO rats are hyperactive and have egocentric and allocentric navigation deficits but showed sparing of conditioned contextual and novel object recognition memory. Here we compared gKO and cKO rats controlling for litter effects. Both gKO and cKO rats were hyperactive and were impaired in egocentric navigation in the Cincinnati water maze (CWM) with deficits greater in gKO rats. The gKO rats were impaired in allocentric navigation in the Morris water maze (MWM) whereas cKO rats were only slightly affected compared with WT, cre, and floxed rats. Striatal tyrosine hydroxylase and dopamine D1 receptors were not significantly different in either model, nor were NMDA-NR1 or NMDA-NR2 in the hippocampus. We previously showed, however, that dopamine is released more rapidly in the striatum of gKO rats by fast-scan cyclic voltammetry. The cKO model shows an important role of catecholamines in the phenotype of LPHN3 disruption and add evidence that this synaptic protein plays a role in neuroplasticity that are consistent with ADHD.

## Introduction

Latrophilin-3 (LPHN3) is an adhesion G-protein coupled receptor (GPCR) located in neuronal terminals where it bridges the synaptic cleft and complexes with FLRT3, Teneurins, and/or UNC5. Variants of *LPHN3* are associated with attention deficit hyperactivity disorder (ADHD) (Arcos-Burgos et al., 2010; Domene et al., 2011a; Gomez-Sanchez et al., 2016; Huang et al., 2018). *LPHN3* variants have been found in those with ADHD where they are associated with combined type ADHD and being responsive to psychostimulants. LPHN3 is highly expressed in brain regions associated with ADHD, including the prefrontal cortex, striatum, and hippocampus (Arcos-Burgos and Muenke, 2010).

Dopamine (DA) is implicated in all models of *Lphn3* deletion (KO) or knockdown (KD), including in *Drosophila melanogaster*, zebrafish, and mice. The KD of *Lphn3* in *Drosophila* using pan-neuronal drivers resulted in persistent hyperactivity that methylphenidate (MPH; 0.5 and 1.0 mg/mL) attenuated (van der Voet et al., 2016).

Zebrafish KD of the orthologue, *lphn3.1*, also resulted in hyperactivity and impulsivity and DA dysregulation (Lange et al., 2012a). *Lphn3.1* morphant hyperactivity was blunted by methylphenidate and duloxetine, consistent with a role for DA in LPHN3- related hyperactivity (Lange et al., 2012b; Lange et al., 2018). A mouse KO of *Lphn3* resulted in short-term (30 min) hyperactivity and increased whole brain DA and serotonin (5-HT) and a heightened locomotor response to cocaine (Wallis et al., 2012; Orsini et al., 2016). They also showed increases in mRNA expression of *Slc6a4* (5-HT transporter), *5-ht2a*, *Drd4*, *Dat1* (DA transporter), neural cell adhesion molecule (*Ncam*), nuclear receptor related 1 (*Nurr1*), and tyrosine hydroxylase (*Th*) [TH (Wallis et al., 2012)]. In the dorsal striatum *Lphn3* KO mice also had increased DA and 5-HT levels compared with WT mice (Wallis et al., 2012).

*Lphn3* KO rats exhibit egocentric navigation deficits in the Cincinnati water maze (CWM) and spatial navigation and cognitive flexibility deficits in the Morris water maze (MWM) (Regan et al., 2021b) along with increased striatal TH, amino aromatic decarboxylase (AADC), and DAT levels (Regan et al., 2019). These data may indicate increased DA turnover resulting in compensatory decreased dopamine D1 receptor (DRD1) and DAARP-32 levels. This idea is consistent with data that gKO rats had increased frequency and amount of striatal DA release by fast-scan cyclic voltammetry (Regan et al., 2020).

Here we report the creation of a floxed *Lphn3* rat and the phenotype when crossed with *Th-cre* rats to generate an *Lphn3-Th-cre* (conditional or cKO) rat. We then compared the cKO with gKO rats.

## Methods

### **Creation of the** *Lphn3-TH-Cre* **line of rats**

*Lphn3^fl/fl^* Sprague Dawley (SD/CD-IGS, strain 001, Charles River, Raleigh, NC) rats were generated through Cincinnati Children’s Transgenic Animal and Genome Editing Core (RRID:SCR_022642) using CRISPR/Cas9 for insertion of LoxP sites around exon 3. *Th-Cre*^HOM-KI RAT,SD,MALE^ (HsdSage:SD-TH^tm1(IRES-Cre)Sage^) rats were purchased from SAGE Labs, Inc. (St. Louis, MO; acquired by Enrivo, Indianapolis, IN). *Lphn3^-/-^ (*knockout) Sprague Dawley rats were generated using CRISPR/Cas9 to delete exon 3 through the Genome Editing Core as described (Regan et al., 2019). Figure 1 shows a schematic of how the conditional *Lphn3-Th-Cre* line was created from the *Lphn3^fl/fl^* and *Th-Cre* lines.

**Figure 1:**
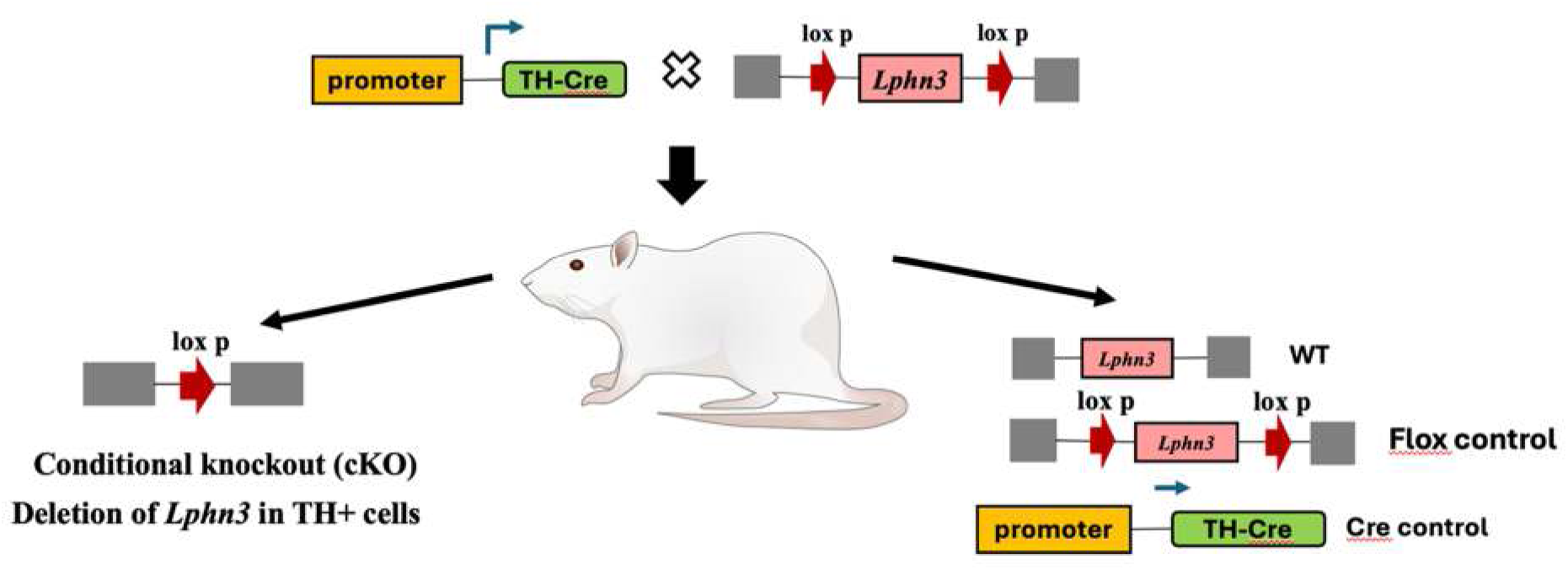
Schematic representation of how the *Lphn3-Th-Cre* conditional KO line of Sprague Dawley rats was created.

### Animal Husbandry

Rats were housed in polysulfone cages in a pathogen free vivarium using the Modular Animal Caging System with HEPA filtered air at 30 air changes/h (Alternative Design, Siloam Spring, AR). Water was reverse osmosis filtered/UV sterilized (SE Lab Group, Napa, CA) and provided ad libitum. Each cage had ad libitum NIH-07 rat chow, woodchip bedding, and a stainless steel enclosure for enrichment (Vorhees et al., 2011). Rats were on a 14 h light/10 h dark cycle (lights on at 600 h). For the gKO, male *Lphn3^+/-^* x multiparous female *Lphn3^+/-^* rats were cohabitated in cages with wire grid floors and checked for sperm plugs. For cKO rats, male *Lphn3^fl/fl^::TH ^IREScre+/-^* x multiparous female *Lphn3^fl/fl^::TH ^IREScre+/-^* were cohabitated in caged with wire grid floors and checked for sperm plugs. Finding a plug was designated embryonic day zero (E0). Birth was E22 (postnatal day zero (P0)). Ear punches were collected on P7 for genotyping. The three primers were used for the gKO rats: 1. AAAGGGTCATAGCATCCGGC, 2. CTAACGTGGCTTTTTGTCTTCT, and 3.

CTCGACAGACAGTGTGGAT. HotStarTaq Master Mix kit (Qiagen Hilden, Germany) was used with DMSO added. Thermocycler parameters were: 1) 94 °C 3 min, 2) 94 °C 3 min, 3) 61.5 °C 30 s, 4) 72 °C 1 min. Steps 2-4 were repeated 15 times and followed by 5) 94 °C 30 s, 6) 59.2 °C 30 s, 7) 72 °C for 1 min, with steps 5-7 repeated 20 times followed by the final steps of 8) 72 °C 10 min, and 9) 4 °C until removed from the machine. The product was run on a 3% agarose gel with ethidium bromide staining. The WT band appears at ∼320 bp and KO band at ∼452 bp.

Ear punches were digested in 200 µL of 50 mM Trizma Base/0.05 % Triton-x100, pH 8.0 and 2 μL proteinase K solution (20 mg/kg Denville Scientific, Metuchen, NJ).

DNA was extracted keeping tissue at 55 °C for 24 h followed by 95 °C for 10 min. Three PCR reactions were used to genotype cKO rats. For the floxed LPHN3, HotStarTaq + Master Mix kit (Qiagen, Hilden, Germany) was used per manufacturer’s recommendation with primers for the 3’ LoxP site: 1. GATGCAGGCATGCTTCGTGT, 2. TTAACCTCCGAGGCACATAACT. Thermocycler parameters were: 1) 94 °C 3 min, 2) 94 °C 1 min, 3) 58 °C for 1 min, 4) 72 °C for 90 s, 5) steps 2-4 were repeated 34 more times followed by 6) 72 °C for 6 min, and 7) held at 4 °C until product was run on a 2% agarose gel with ethidium bromide. The WT band occurs at ∼431 bp and the 3’ LoxP site band at ∼477 bp. For the *TH^IREScre+^* band, Choice Taq Blue Master mix (Denville Scientific, Metuchen, NJ) was used per manufacturer’s recommendation with primers: 1. TCCCTCCACAGGAACTATGC, 2. TATAGAGCATGGAGGGCAGG, and for positive control 3. TTACGTCCATCGTGGACAGC, 4. TGGGCTGGGTGTTAGCCTTA.

Thermocycler parameters were: 1) 95 °C 5 min, 2) 95 °C 30 s, 3) 60 °C 45 s,4) 72 °C 90 s, 5) 95 °C 30 s, 6) 60 °C 45 s, 7) 72 °C 90 s, 8) steps 5-7 repeated 18 times with step 6 as a touchdown PCR ramping down 0.5 °C/cycle, 9) 94 °C 30 s, 10) 50 °C 45 s, 11) 72 °C 90 s, 12) repeated steps 9-11 13 times, 13) 72 °C 10 min, and 14) held at 10 °C until the product was run on a 2% agarose gel with ethidium bromide for staining. The *TH^IREScre+^* band occurs at ∼400 bp and the positive control band at ∼184 bp. For the *TH^IREScre-^* band, similar methods were used, except using primers: 1.

GAACCTGATGGACATGTTCAGG, 2. AGTGCGTTCGAACGCTAGAGCCTGT, and the same primers for the positive control. The *TH^IREScre-^* band occurs at ∼324 bp.

Litters were weaned on P28, and offspring housed 2/cage/sex. A total of 180 rats were used (33 *Lphn3^fl/fl^::TH ^IREScre+/+^* (cKO), 28 *Lphn3^fl/fl^::TH ^IREScre-/-^* (Flox), 29 *Lphn3^+/+^::TH ^IREScre+/+^* (Cre)*, and* 32 *Lphn3^+/+^::TH ^IREScre-/-^* (cWT), 29 gKO and 29 WT). After behavior, brains were removed and dissected for RT-PCR. A separate cohort of 4 Cre and *Lphn3* cKO rats were used for immunohistochemistry. To help control for litter effects, only one rat per genotype per sex was tested per litter. Rats were tested in our institution’s Animal Behavioral Shared facility (RRID:SCR_022621) by personnel blind to genotype. All protocols were approved by the Cincinnati Children’s Institutional Animal Care and Use Committee and conformed with relevant guidelines and regulations.

### Behavior

For home-cage testing at P35, rats from 47 litters were used for the conditional groups (17 cKO females, 16 cKO males, 17 cWT females, 14 cWT males, 15 cre females, 13 cre males, 14 floxed females, 13 floxed males). For home-cage testing at P50, rats from 49 litters for the conditional group were used. For the global KO groups, rats came from 20 gKO litters were used (15 gWT females, 14 gWT males, 15 gKO females, and 14 gKO males) at both P35 and P50.

After P50 behavioral tests were performed in the following order: acoustic and tactile startle response habituation (ASR-TSR), acoustic prepulse inhibition of acoustic and tactile startle, light prepulse inhibition of acoustic and tactile startle, straight swim channel, Cincinnati water maze (CWM), Morris water maze (MWM), radial water maze (RWM), and mirror image CWM. A week after the last test, the hippocampus, caudate-putamen, nucleus accumbens, prefrontal cortex, and cerebellum were collected and frozen. Behavioral equipment was cleaned between rats with Process NPD (STERIS Life Sciences, Mentor, OH) an EPA approved, non-toxic denaturing, antibacterial, antiviral cleaning agent.

### Home-cage

Rats were housed singly in standard cages during this test (Tang et al., 2002). Each cage rested in a metal frame with infrared photodetectors spaced 5 cm apart along the X and Y axes. The frame was positioned 2 cm above the cage floor. Data were collected for 72 h at P35 and P50 (PAS System, San Diego Instruments, San Diego, CA). Because of an error, only movement in the central region of the home-cage was analyzed.

### Acoustic/Tactile Startle Habituation

ASR and TSR habituation were assessed in SR-LAB apparatus (San Diego Instruments, San Diego, CA). Rats were tested on 2 successive days. On each day, rats were placed in acrylic cylindrical holders mounted on a platform with a piezoelectric accelerometer attached to the underside. This assembly was placed inside a sound-attenuated cabinet containing a house light and fan. Background noise was 55 dB. Each session consisted of a 5 min acclimation period followed by 100 trials. The acoustic pulse was a 20 ms, 120 dB mixed frequency white noise burst (rise time 1.5 ms). The tactile pulse was a 60-psi air-puff to the dorsal surface from a tube mounted through a slot in the top of the animal holder. Movement detected at the start of each trial was subtracted from the maximum startle amplitude (V_max_) to eliminate non-startle movement artifacts. A V_max_ < 95 mV was considered a nonresponse for that trial and the data were excluded.

### Acoustic Prepulse Inhibition (PPI) of Acoustic and Tactile Startle

PPI was tested in SR-LAB test chambers as above (San Diego Instruments, San Diego, CA). Rats were given 100 trials in a 10 x 10 Latin square sequence of 100 trials repeated 2 times with trial types being pulse alone (acoustic or tactile) or prepulse + pulse. Blocks of 5 were given for acoustic and tactile trials. Acoustic prepulses were 59, 70, 80, or 93 dB. Prepulses preceded pulses by 70 ms from prepulse onset to pulse onset; hence, the signal gap was 50 ms. The pulses were the same as above. Trials of the same type were averaged for analysis. V_max_ was the dependent measure. If the unmodified V_max_ < 95 mV for a block of 5 trials, then data for all trials of that type (ASR or TSR) within that block were removed from the analyses.

### Light Prepulse Inhibition (PPI) of Acoustic Startle

The acoustic and tactile pulses were as above. The light prepulse was from ceiling mounted LEDs (∼1100 lux). Each rat received 200 trials of five types: no prepulse (acoustic or tactile pulse) or light prepulse given 30, 70, 100, or 400 ms before the pulse. Trials were presented in 10 x 10 Latin square sequence. All other measures were as with the acoustic PPI.

### Straight Channel

Rats were given 4 trials to escape a 244 cm long x 15 cm wide x 50 cm high straight channel filled halfway with room temperature water (20 °C). Latency to reach a platform from the opposite end was recorded. The test acclimates rats to swimming and provides practice escaping on a hidden platform. Latencies were used to compare groups for equivalent swimming performance and motivation.

### Cincinnati Water Maze (CWM)

The CWM assesses egocentric, route-based learning and memory (Vorhees and Williams, 2016). The maze consists of 10 T-shaped cul-de-sacs that branch from a central channel extending from the start to the goal where a submerged escape platform is located. To exclude distal cues, testing was conducted in the dark using only infrared light. A video camera sensitive to infrared was mounted above the maze and connected to a monitor in an adjoining room where the experimenter monitored performance. Rats were acclimated to the dark for at least 5 min prior to testing. Rats were monitored and scored for latency to reach the goal and errors. An error was a head and shoulder entry into the stem or arm of a T-section. Rats that reached the 5 min limit without completing the task had errors adjusted to the rat that made the most errors. There were two trials per day for 18 days. If a rat did not find the platform on trial-1 of a given day, it was placed in a holding cage for at least 5 min before trial-2, otherwise trials were given back-to-back. We used two versions of this maze (Figure 2), the original and its mirror image.

**Figure 2:**
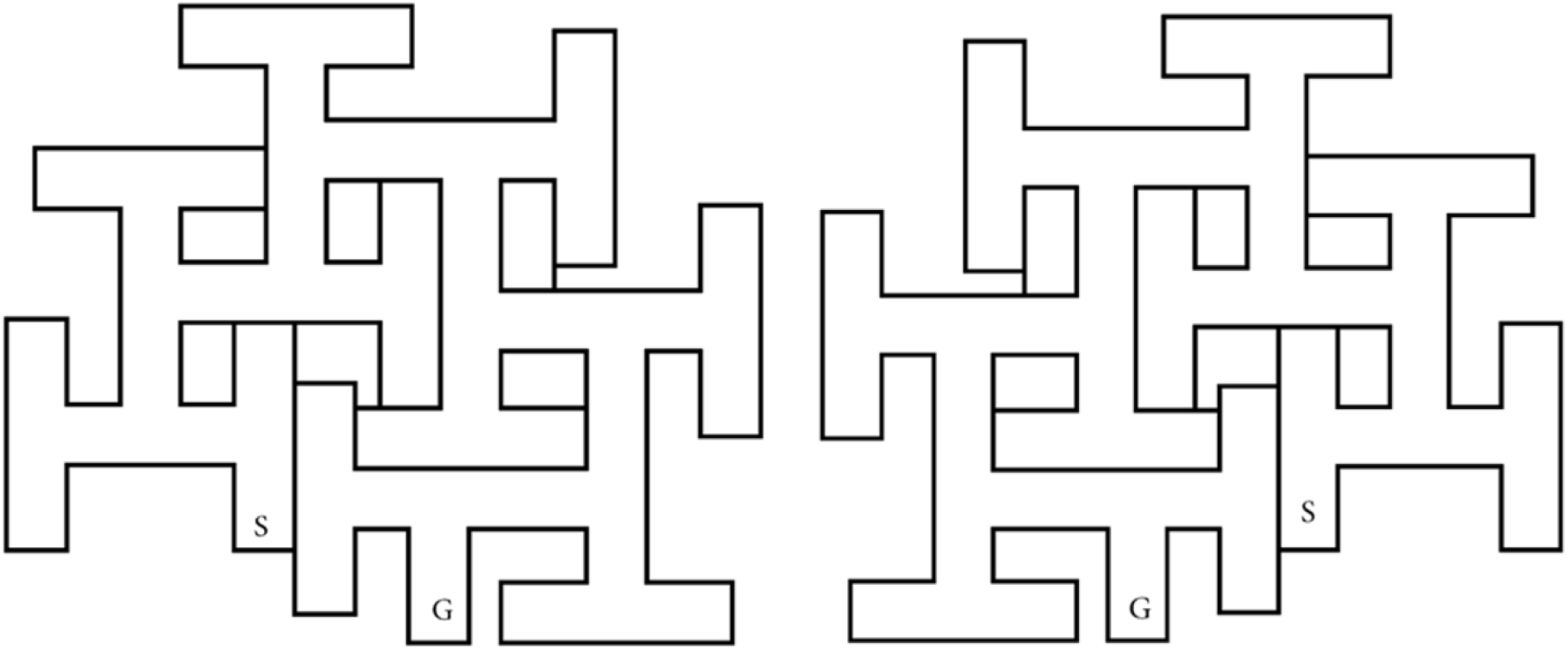
Schematic of the Cincinnati water maze (CWM). Left: The standard configuration; Right: Mirror image of the standard configuration.

### Morris Water Maze (MWM)

To assess allocentric learning and memory, rats were tested in a MWM (Vorhees and Williams, 2006; Vorhees and Williams, 2014). The tank was constructed of black laminated polyethylene and is 244 cm in diameter and 51 cm deep, filled halfway with room temperature water. Curtains were mounted on tracks on the ceiling that could be opened or closed to expose or hide distal cues (geometric shapes and posters). Rats were tested in 4 phases: acquisition, reversal, shift, and cued-random. The first three phases consisted of 4 trials per day for 6 days; the last phase was 2 days. Two probe trials were given for acquisition, reversal, and shift: one on day-3 before the platform trials for the day were given and one 24 h after the last day of platform trials (day-7).

Probe trials lasted 45 s. For learning trials, the time limit was 2 min. Rats were tested in rotation, i.e., all rats in a subgroup completed trial-1 before being given trial-2, etc. If a rat did not find the platform within 2 min, it was lifted out and placed on the platform for 5 s. During platform trials, the platform was 2 cm below the surface and positioned halfway between the center and wall of the tank. For acquisition a 10 cm diameter platform was placed in the SW quadrant. Rats started from one of two cardinal and two ordinal positions around the perimeter distal to the goal. During reversal, a 7 cm diameter platform was positioned in the NE quadrant and start positions adjusted accordingly. In the shift phase, a 5 cm diameter platform was placed in the NW quadrant and start positions adjusted. A camera mounted above the maze was synchronized to a computer with video tracking software (AnyMaze, Stoelting Co., Wood Dale, IL). For learning trials, dependent variables were latency, path efficiency, and swim speed. Dependent measures on probe trials were average distance to the former platform site, entries, and swim speed.

The fourth phase was cued-random. Curtains were closed around the pool to conceal distal cues to assess proximal cue learning. A plastic ball protruded 10 cm above the water that was affixed to a metal rod mounted at the center of the platform. Rats were given 4 trials/day for 2 days. Positions of the platform and start locations were pseudo-randomized on every trial to prevent use of any residual distal cues not masked by the curtains. Latency was recorded.

The maze tests explicit/spatial/allocentric learning and reference memory (probe trials). Acquisition assesses spatial learning, reversal assesses cognitive flexibility, and shift assesses cognitive flexibility with greater retroactive interference. The cued- random phase tests visually guided egocentric learning.

### Radial Water Maze (RWM)

To assess working memory, rats were tested in an 8-arm RWM. The tank was 208 cm diameter x 56 cm deep made of black polyethylene and filled with water to a depth of 32 cm. Posters were placed on the walls to serve as distal cues. The start was from arm-0; the other arms were numbered clockwise, 1-7; each contained a submerged platform. There were 7 trials/day for 2 days. Working memory errors were counted when a rat reentered an arm it previously entered, start errors were counted when a rat reentered the start arm; total errors were the sum of working and start errors. Rats were placed at the start, and time to reach a platform and errors recorded. Once a platform was reached, the rat was removed after 5-10 s and placed in a holding cage for 30 s while the platform they found was removed, leaving 6 platforms for trial-2. This continued until all platforms were found. The trial limit was 2 min.

### Cincinnati Water Maze mirror image

Rats were tested in a CWM that was a mirror-image of the first one. The same procedures were used as above (2 trials/day for 10 days).

### Viral Injections

A separate group of 4 male cre rats were anesthetized with 2–4% isoflurane (IsoThesia; Butler Animal Health Supply, Dublin, OH) with continuous administration via a nose cone throughout surgery. Adult Cre rats were placed in a motorized, computer-controlled stereotaxic apparatus (StereoDrive, Stoelting Co., Wood Dale, IL) and given bilateral injections of pAAV-FLEX-tdTomato 28306 (Addgene, Watertown, MA) using a 26 gauge 10 μL Hamilton Gastight syringe (Reno, NV). pAAV-FLEX-tdTomato was a gift from Edward Boyden (28306-AAV9; http://n2t.net/addgene:28306 ; RRID:Addgene_28306). Virus titers were 3.8 x 10^13^ gc/mL for AAV9. Coordinates were based on the Paxinos and Watson brain atlas (Paxinos and Watson, 2006). For the dorsal lateral striatum (DLS), a volume of 3 μL was injected over 9 min (from bregma: AP: +3.2 mm; ML: ± 2.0 mm; from skull: DV: −6.5 mm), with the needle left in place for 5 min following injection.

### Immunohistochemistry

Four weeks after surgery, rats were perfused transcardially with 4% paraformaldehyde, brains dissected, postfixed, and placed in sucrose overnight. Brains were sectioned (30 μm) on a cryostat, and free-floating sections processed for TH immunohistochemistry, using sheep anti-TH primary antibody (ab113, diluted 1:100; AbCam, Cambridge, MA) with anti-sheep secondary antibody at 1:500 with DAPI counterstain. Images were taken using a Nikon confocal microscope (Microscopy Imaging Core); all images are 20x magnification.

### Quantitative PCR

Quantitative PCR (qPCR) was used to analyze gene expression of *Lphn3* in the caudate-putamen (CPu) from 6 gKO, 6 cKO, 6 gWT, and 6 cWT male and female rats. RNA was isolated using RNAqueous-Micro (ThermoFisher Scientific) following the manufacturer’s instructions. RNA was isolated using 1 mL of TRIzol for every 50-100 mg of tissue and quantified by Nanodrop (Thermo Scientific). Reverse transcription reactions were performed using 4 µL of iScript at room temperature with 1 µg-1 pg of RNA template (Bio-Rad) in a final volume of 20 μL. PCR reactions were carried out as follows: 5 min at 25 °C, 20 min at 46 °C, and 1 min at 95 °C. The qPCR samples contained 160 ng of cDNA, 300 nM of each primer (forward and reverse), and 1x SYBR Green Master Mix (BioRad) in a 20 µL volume. Two 20 µL aliquots of the mix were placed in a 96-well plate and the qPCR was performed on a 7500 Real Time PCR System (Applied Biosystems) using the following conditions: 50 °C for 2 min, 95 °C for 10 min, 50 cycles at 95 °C for 15 s, and 60 °C for 1 min. Primers were synthesized by Integrated DNA Technologies (Coralville, IA) and selected based on primer efficiency of 95-100%. Rat primer sequences were as reported (Regan et al., 2019). Ct values were determined by the SDS 2.4 software with a threshold set at 0.5. The average Ct values from assayed duplicates were calculated and averaged. Changes in mRNA were measured with the ΔΔCt method (Livak and Schmittgen, 2001) using actin as the reference and *Lphn3* WT samples as calibrator.

### Data Analyses

Data were analyzed by generalized linear mixed-effect models using SAS (v9.4, SAS Institute, Cary, NC) with p ≤ 0.05 as the threshold for significance. To control for litter effects and sex within litter only one rat per genotype per sex per litter was used and litter and litter x sex were random factors in the statistical analyses (Golub and Sobin, 2019; Vorhees and Williams, 2019). The gKO and the cKO genotypes were analyzed separately. Two factor mixed linear model ANOVAs were used with between-subject factors of genotype and sex and data presented as least square mean (LS Means) ± standard error . Variance-covariance matrices of best fit were used, either autoregressive (AR) or AR moving average together with Kenward-Roger first order degrees of freedom. The repeated measure was time for home-cage, trials for straight channel, trial blocks for startle, and test day for mazes. Significant interactions were analyzed using slice-effect ANOVAs in SAS (simple main effects in a single procedure with the overall error term maintained) for genotype along the repeated measure dimension. cWT compared with cKO were made using Dunnett tests.

## RESULTS

### Reproductive Outcomes

**Table** **1** shows basic characteristics of the Conditional line compared with the Global line of rats. There was a significant difference between the two with the average litter size of the Global line smaller than for the Conditional line.

**Table 1:** Reproductive Outcomes^1^

| **Table 1** |  |  |
| --- | --- | --- |
| Reproductive Outcomes<sup>1</sup> |  |  |
| **Pups/litter** |  |  |
| **Average No./litter** | **Conditional**<br>(cKO+cWT+Cre+Flox) | **Global**<br>(gKO+gWT) |
| Litter size | 11.9 ± 0.3 | 10.8 ± 0.3** |
| Males | 6.0 ± 0.2 | 5.7 ± 0.2** |
| Females | 5.9 ± 0.2 | 5.1 ± 0.2** |
| **P7 Body wt. (g)<sup>2</sup>** |  |  |
| Males | 19.27 ± 0.29 | 17.68 ± 0.33** |
| Females | 18.47 ± 0.30 | 17.01 ± 0.33** |
| <sup>1</sup>Mean ± SEM |  |  |
| <sup>2</sup>Body weight (g) prior to genotyping |  |  |
| **P<0.01 Global vs. Conditional genotypes |  |  |

### Body Weight

There were differences between the two genetic lines in terms of growth as reflected in body weight (Fig. 3). For the Global line there was a significant effect of genotype [(1,124) = 90.14, p<0.0001)], genotype x sex [(1,123) = 4.62, p = 0.034)], and a nearly significant genotype x sex x age [(16,1216) = 1.64, p = 0.052)]. gKO rats weighed significantly less by P21 compared with gWT rats, a difference that persisted into adulthood (Fig. 3E). For the Conditional line, there was a significant effect of genotype [(3,222) = 0.038). The cKO rats never differed significantly from cWT, cre, or floxed rats (Fig. 1), however the floxed rats weighed significantly less as adults than did the cWT rats (p = 0.026).

**Figure 3:**
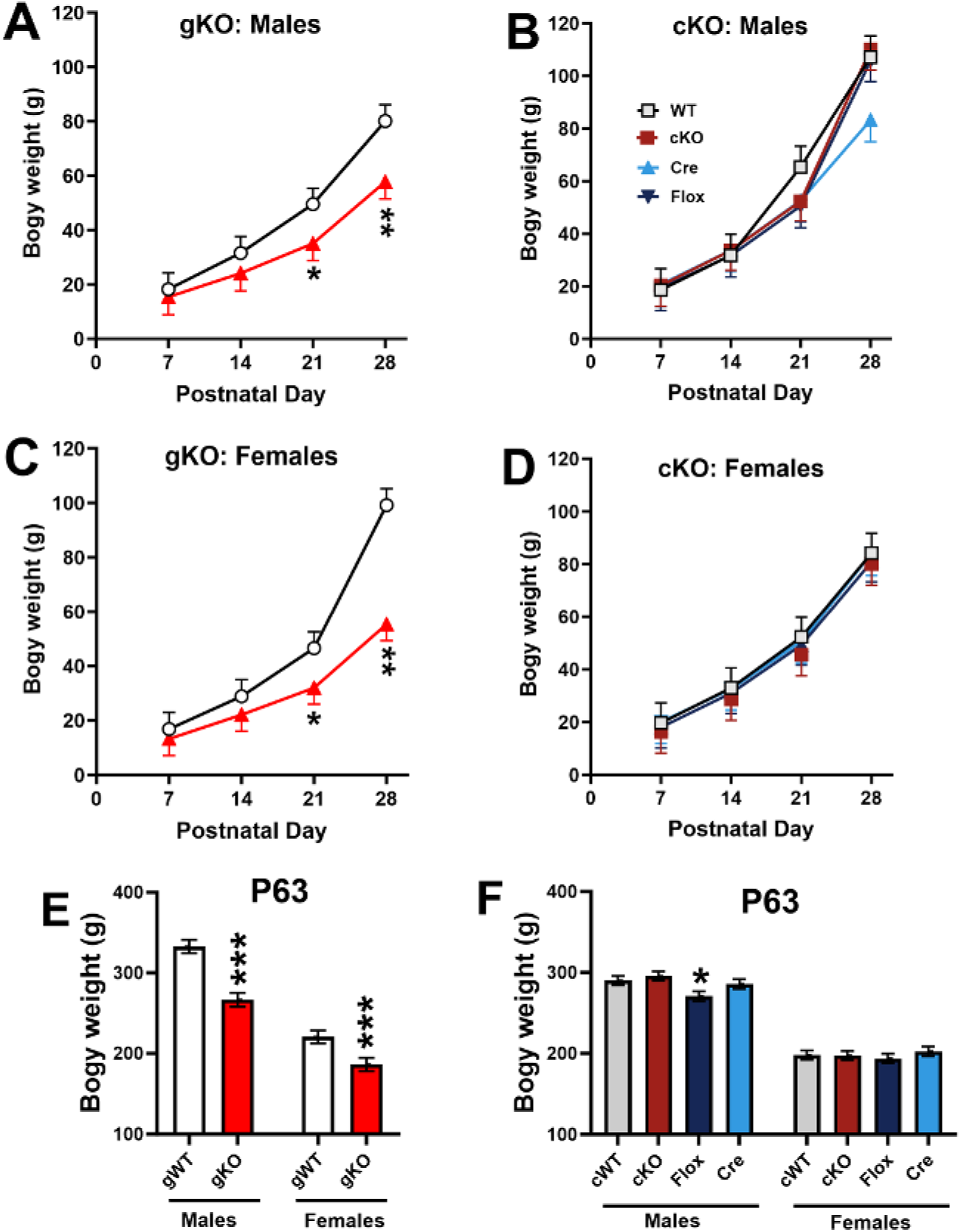
Preweaning and adult body weight. **A**, male gKO and gWT body weights (g). **B,** male cKO, cWT, Cre, and Flox body weights. **C,** female gKO and gWT body weights. **D,** female cKO, cWT, Cre, and Flox body weights. *p<0.05, **p<0.01 vs. WT. Group sizes: Preweaning gKO male = 35-36, gWT male = 36-38; gKO female = 35-36, gWT female = 34-36. Adult (P63) male gKO = 15; male gWT = 16; female gKO = 17, female gWT = 18. Preweaning Male cKO = 32-35, cWT = 30-31, Cre = 29-31, Floxed = 28-32; female cKO = 30-31, cWT = 30-32, Cre = 28-31, Floxed = 32-34; Adult (P63) Male cKO = 21, cWT = 32, Cre = 14, Floxed = 15; Female cKO = 20, cWT = 21, Cre = 16, Floxed = 19.

### Home-cage

At P35, the main effect of genotype was not significant in the gKO but there was a significant genotype x interval [F(35,1607) = 1.64, p = 0.0106], Fig. 4A, however there were no significant genotype or interactions with genotype in the cKO (Fig. 4B)

**Figure 4.**
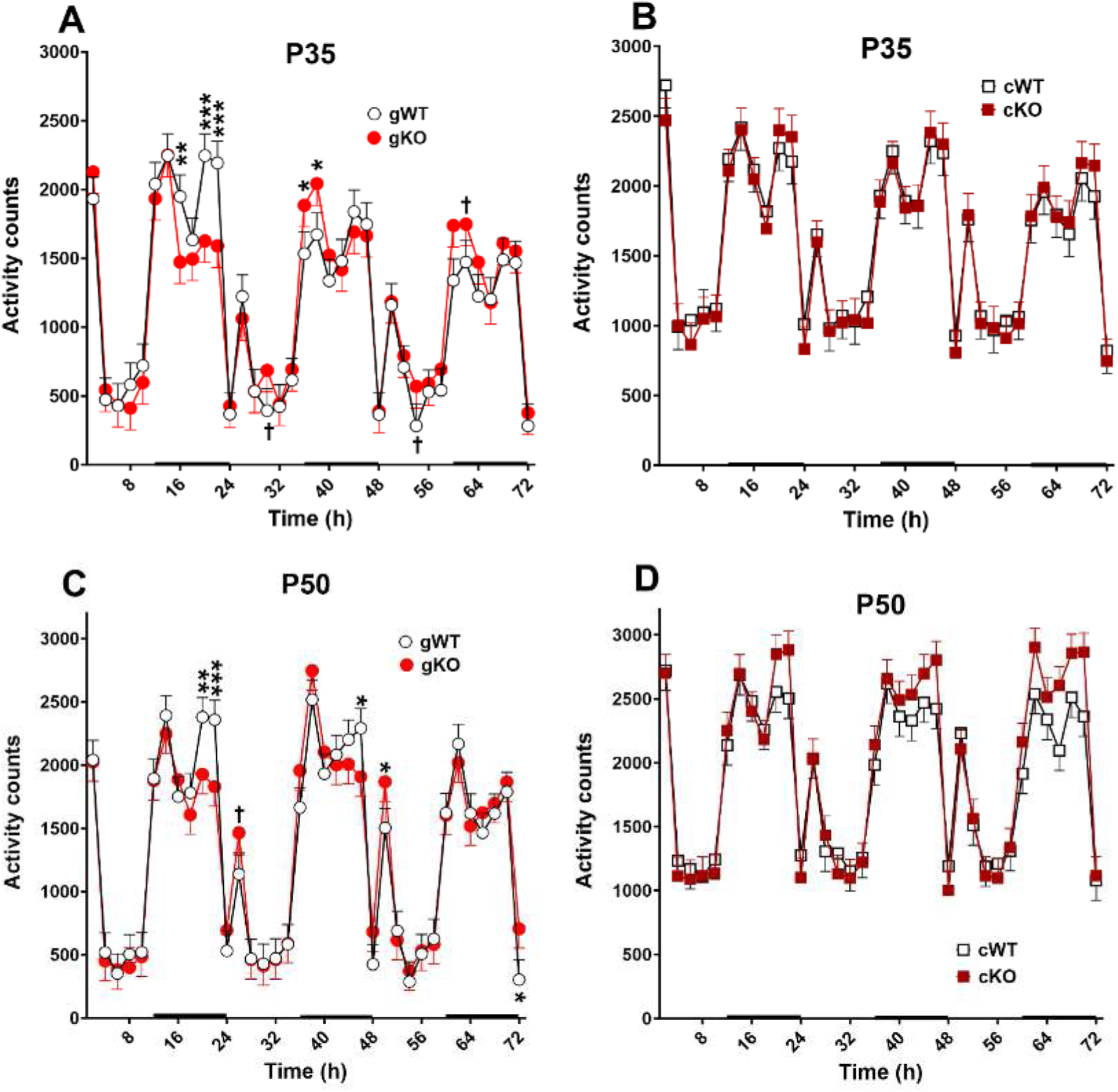
Home-cage: Home-cage locomotor activity. Rats were tested 72 h, starting 8 h before the dark cycle on a 14 h light, 10 h dark cycle. **A,** gKO vs. gWT at P35. **B,** cKO vs. cWT at P35. **C,** gKO vs. gWT at P50, **D**, cKO vs. cWT at P50. There was no genotype × sex interaction, therefore, male and female data were combined. Group sizes: P35: cKO: 17 F, 16 M; cWT: 17 F, 14 M; gWT: 15 F, 14 M, gKO: 15 F, 14 M. P50: cKO: 17 F, 16 M; cWT: 17 F, 14 M; gWT: 15 F, 14 M, gKO: 15 F, 14 M. †P<0.10, *P<0.05, ***P<0.001 vs WT.

At P50, there was no main effect of genotype for the gKO rats but there was a genotype x interval interaction [F(35, 1666) = 1.50, p = 0.0304]. Slice comparisons at each interval are shown in Fig. 4C. At P50, genotype was not significant nor was the genotype x interval, but there was a significant genotype x sex interaction [F(3,137) = 3.23, p= 0.0243] in which female cKO rats were more active than female cWT with no differences among males but a significant difference among females [F(3,136.6) = 2.88, p = 0.0384]. The cKO and Cre groups differed from cWT (p = 0.0193, and 0.0114, respectively, Mean ± SEM: cWT: 2144.2 ± 106.8, cKO: 1919.8 ± 116.6, Flox: 2039.4 ± 115.5, Cre: 1890.7 ± 112.7).

### Acoustic and Tactile Startle Habituation

For acoustic startle, gKO rats had increased responses compared with gWT rats [F(1, 26.5)= 13.74, p<0.001] (Fig. 5A). Responses, regardless of genotype, were greater on day-2 than on day-1 [Day: F(1, 43) = 5.86, p<0.01]. For cKO rats, there was a main effect of genotype [F(3, 82.7) = 4.00, p<0.05]. Flox rats had increased startle compared with cre, cWT, and cKO rats.

**Figure 5:**
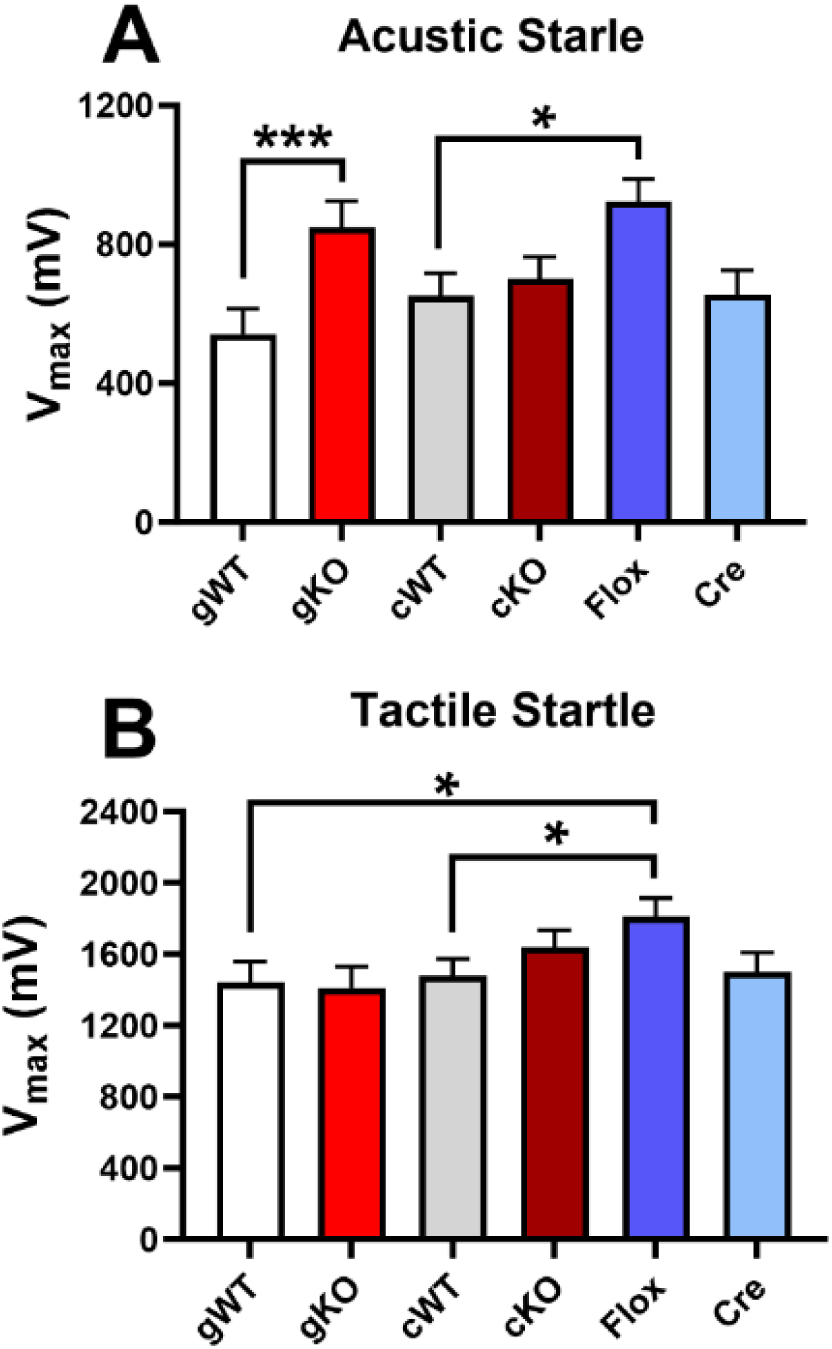
Acoustic and Tactile Startle Habituation: A, Acoustic startle. **B,** Tactile startle. Group sizes: cKO: 17 F, 16 M; cWT: 17 F, 14 M; Cre: 15 F, 13 M; Floxed: 14 F, 13 M; gWT: 15 F, 14 M, gKO: 15 F, 14 M.*P<0.05, ***P<0.001.

For tactile startle, gKO rats were not different from gWT rats. For cKO rats, there was a main effect of genotype [F(3, 78.2)= 2.85, p<0.05], with flox rats having increased startle compared with cWT rats (Fig. 5B). There was no effect of day or genotype x day. Males had greater responses compared with females [F(1, 77.8) = 10.66, p<0.01].

### Acoustic and Tactile PPI

For acoustic PPI of the acoustic startle response, there was no main effect of genotype for gKO rats but there was an interaction of genotype x prepulse [F(4, 110) = 3, p <0.05], where gKO rats had an increased response at PP-59 and a trend at in PP- 70 (Fig. 4A) compared with gWT rats. cKO rats had a main effect of genotype [F(3, 80.8) = 2.71, p <0.05]. Post hoc analyses showed that flox rats had increased responses compared with cre and cKO rats (Fig. 4B). There was no genotype x prepulse interaction.

For acoustic PPI of the tactile startle response, gKO rats had increased responses compared with gWT rats [F(1, 17.6) = 5.15, p<0.05] (Fig. 4D). There was a trend for a genotype x prepulse interaction [F(4,102)= 2,15, p=0.08], Fig. 4C). The cKO rats had increased responses compared with cWT and cre rats [F(3, 83.7) = 5.09, p<0.01] (Fig. 3D). There was no genotype x prepulse interaction.

### Acoustic and tactile startle with light PPI

For acoustic startle with light prepulse, neither gKO nor cKO rats had a genotype effect or genotype-related interactions (Fig. 6E,F).

**Figure 6:**
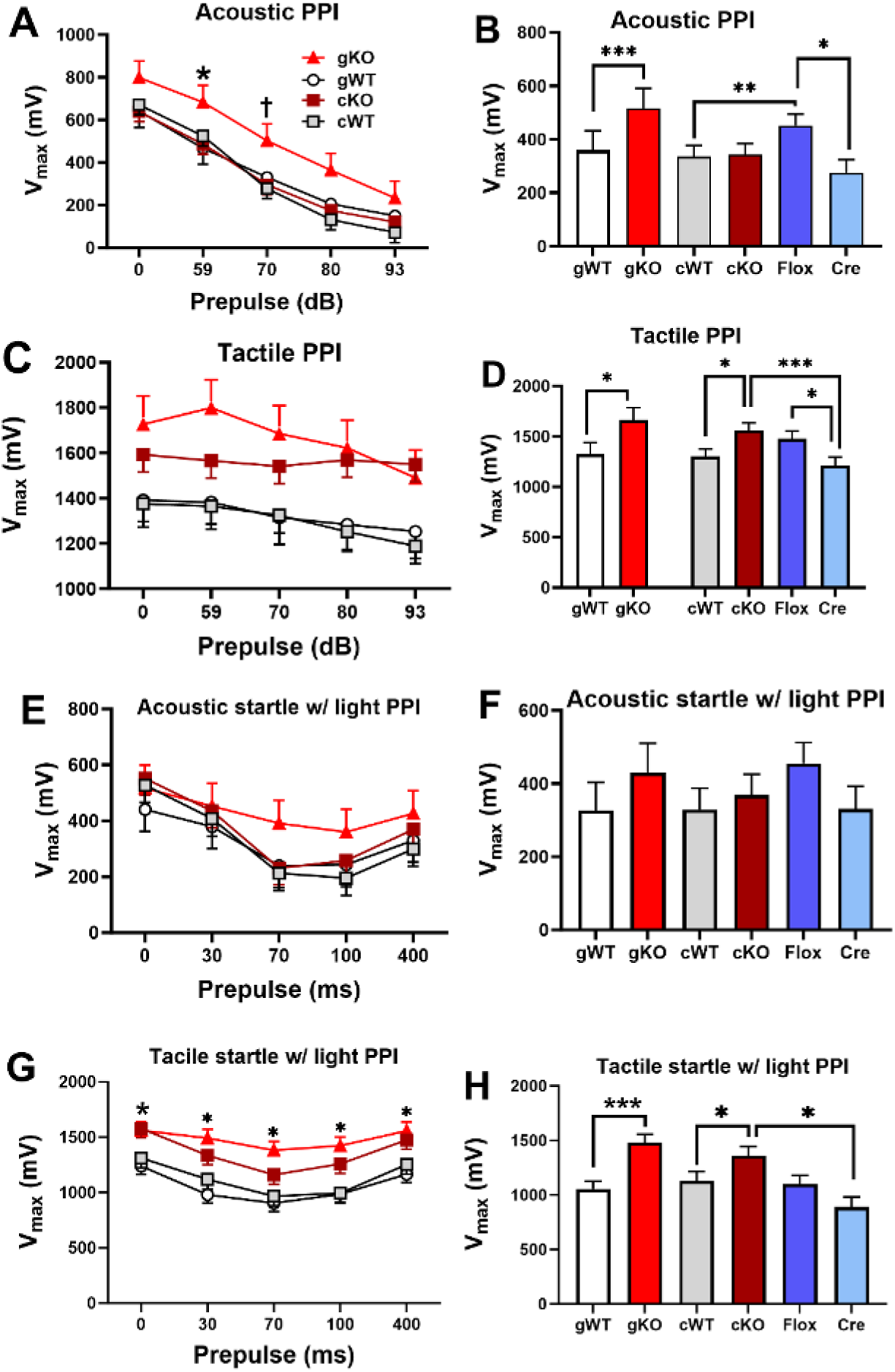
Acoustic and Tactile Startle with acoustic or light prepulses (PPI): A,B, Acoustic startle with acoustic prepulse. **C,D,** Tactile startle with acoustic prepulse. **E,F**, Acoustic startle with light prepulse. **G,H**, Tactile startle with light prepulse. Group sizes: cKO: 17 F, 16 M; cWT: 17 F, 14 M; Cre: 15 F, 13 M; Floxed: 14 F, 13 M; gWT: 15 F, 14 M, gKO: 15 F, 14 M.*P<0.05, **P<0.01, ***P<0.001.

For tactile startle with light prepulse, gKO rats had an increased response compared with gWT rats [F(1, 34) = 17.19, p<0.001] (**Fig.** **6G,H**). There was an interaction of genotype x prepulse [F(4, 136) = 3.77, p<0.01] where gKO rats had increased responses at each prepulse level. There was an interaction of sex x genotype [F(1, 34) = 5.86, p<0.05] where gKO males had increased responses compared with gWT males, but this was not significant for females (not shown). cKO rats had increased responses compared with cre, floxed, and cWT rats, and cre rats had decreased responses compared with cWT rats [F(3, 83.7) = 5.09, p<0.01]. There were no genotype-related interactions.

### Straight Channel

There was a main effect of genotype [Genotype: F(1, 68.2) = 15.81, p<0.0001] in which gKO rats took longer to reach the platform than gWT rats (Fig. 7A). There was no genotype effect for cKO rats. There were no genotype-related interactions.

**Figure 7:**
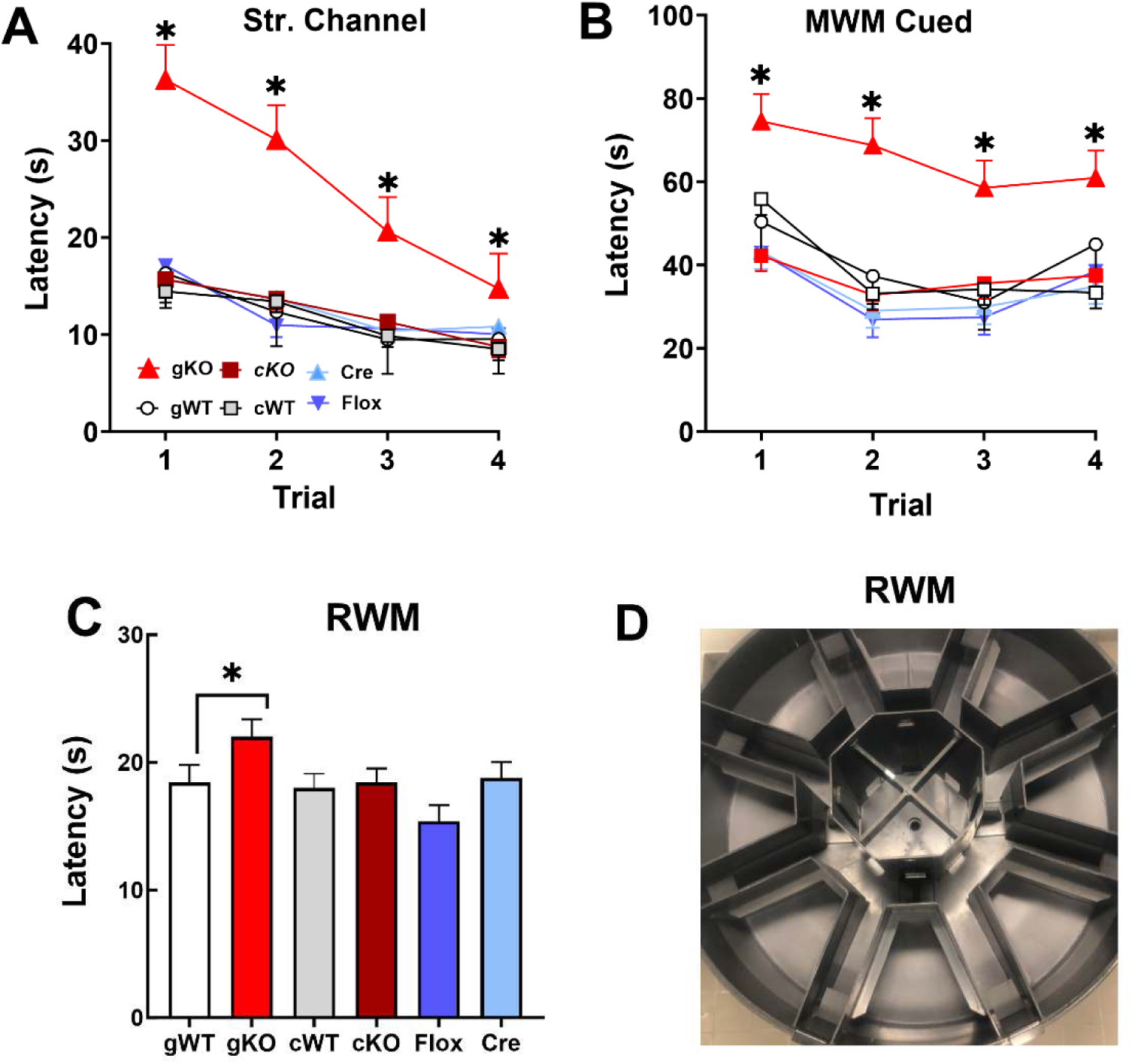
Straight Channel, RWM, MWM-Cued, RWM illustration: A, Straight Channel latency to reach a submerged platform swimming from one end to the other. **B,** MWM Cued (visible platform) trials. **C,** RWM latency to reach an escape platform. **D**, RWM apparatus. **Group sizes:** cKO: 17 F, 16 M; cWT: 17 F, 14 M; Cre: 15 F, 13 M; Floxed: 14 F, 13 M; gWT: 15 F, 14 M, gKO: 15 F, 14 M.*P<0.05 **^#^**P<05 or beyond.

### RWM

The apparatus is shown in Figure 7D. There was a main effect of genotype on latency [F(1,30.6) = 5.42, p=0.0266] in which the gKO rats had increased latencies compared with gWT rats (Fig. 7C). There were no significant effects for the cKO rats.

### CWM

gKO rats had longer latencies to escape than gWT rats [Genotype: F(1,47) = 8.62, p<0.01] (Fig. 8A,B). There was no genotype x day interaction. Males had longer latencies than females [Sex: F(1, 64.5) = 4.63, p < 0.05]; there were no genotype interactions. cKO rats had longer latencies than cWT, cre, or flox rats [Genotype: F(3,123) = 4.74, p<0.01] (**Fig** **8A,B**). There was a trend of genotype x day (p = 0.09). Males had longer latencies than females [F(1,128)= 3.80, p<0.05]; there were no genotype interactions.

**Figure 8:**
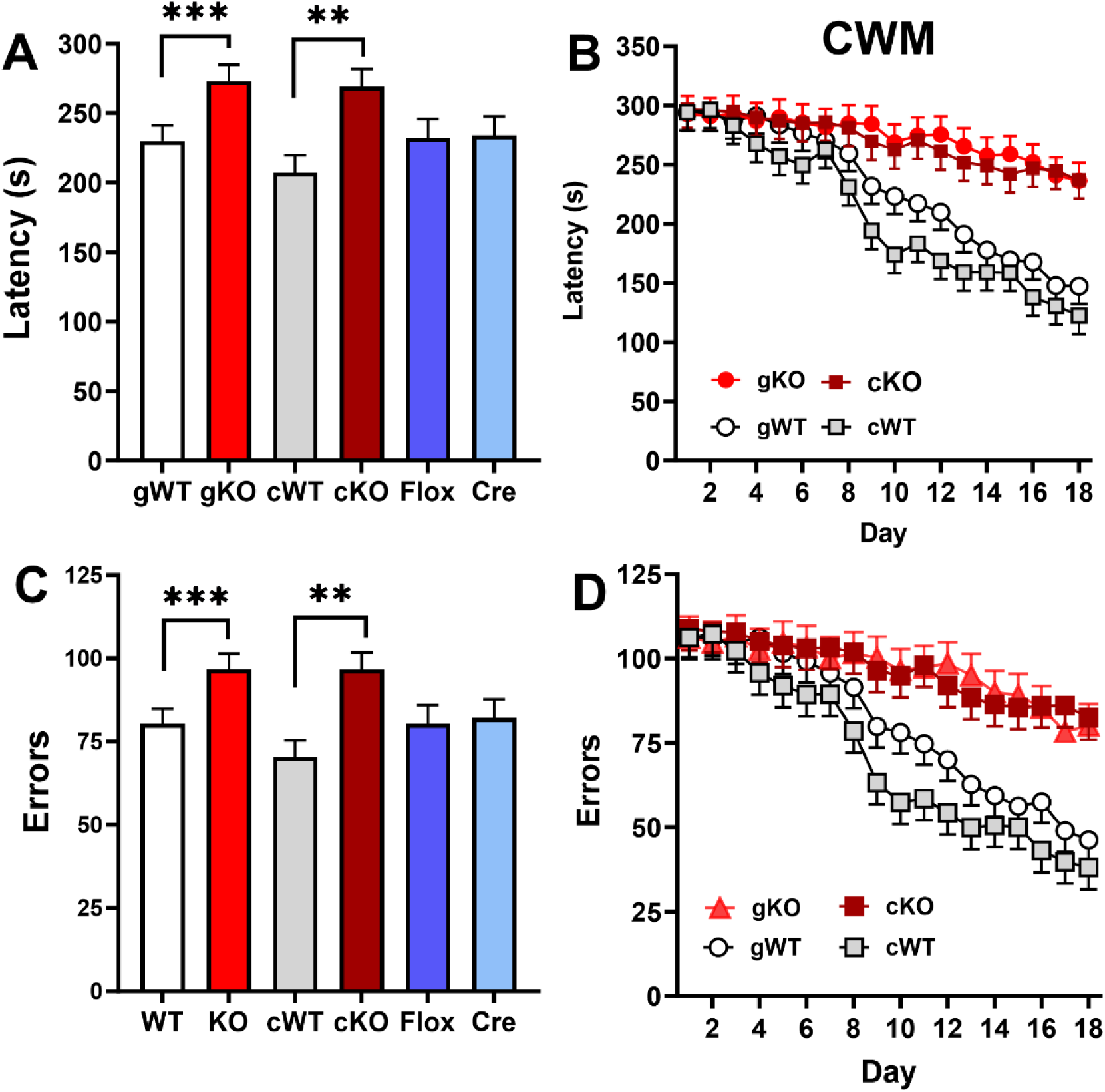
Cincinnati Water Maze: A,B, **g**KO and cKO rats were significantly different from gWT and cWT rats on both latency and errors. **A,** latency main effect; **B,** latency by day (2 trials/day); **C**, main effect for errors; **D**, errors by day. **Group sizes:** cKO: 17 F, 16 M; cWT: 17 F, 14 M; Cre: 15 F, 13 M; Floxed: 14 F, 13 M; gWT: 15 F, 14 M, gKO: 15 F, 14 M.**P<0.01, ***P<0.001.

The gKO rats made more errors than gWT rats [genotype: F(1,47.1) = 7.81, p < 0.01] (Fig. 8C,D). Males had more errors than females [F(1, 64.8) = 4.60, p <0.05].

There were no genotype interactions. cKO rats had increased errors compared with cre, floxed, and cWT rats [genotype: F(3,124) = 4.98, p < 0.01] (Fig. 8C,D). There were no differences between cre, floxed, and cWT rats. There were no genotype interactions. Males showed a trend toward making errors than females (sex: p = 0.06).

### Morris water maze

For acquisition, gKO rats had longer latencies compared with gWT rats [Genotype: F(1, 69.4) = 47.62, p<0.0001] (Fig. 9A). There were no genotype interactions. gKO rats were also less efficient at reaching the platform compared with gWT rats [Genotype: F(1, 58.7) = 71.29, p<0.0001] (Fig. 9B); there were no interactions with genotype. There were no differences in swim speed; mean ± SEM were: gKO, 19.59 ± 0.84 cm/s; gWT, 11.37 ± 0.84 cm/s. cKO rats had longer latencies compared with cWT, floxed, and cre rats [Genotype: F(3, 192) = 6.88, p<0.001] (Fig. 9A). There were no genotype-related interactions. There was a main effect of genotype for cKO rats for path efficiency [F(3, 187) = 2.73, p< 0.05]. The cKO rats had lower path efficiency compared with cWT rats (Fig. 9B). There were no genotype-related interactions. There were no genotype effects on swim speed; mean ± SEM were: cKO, 24.58 ± 0.40 cm/s; cWT, 24.10 ± 0.40 cm/s; flox, 25.34 ± 0.44 cm/s; cre, 24.18 ± 0.43cm/s.

**Figure 9:**
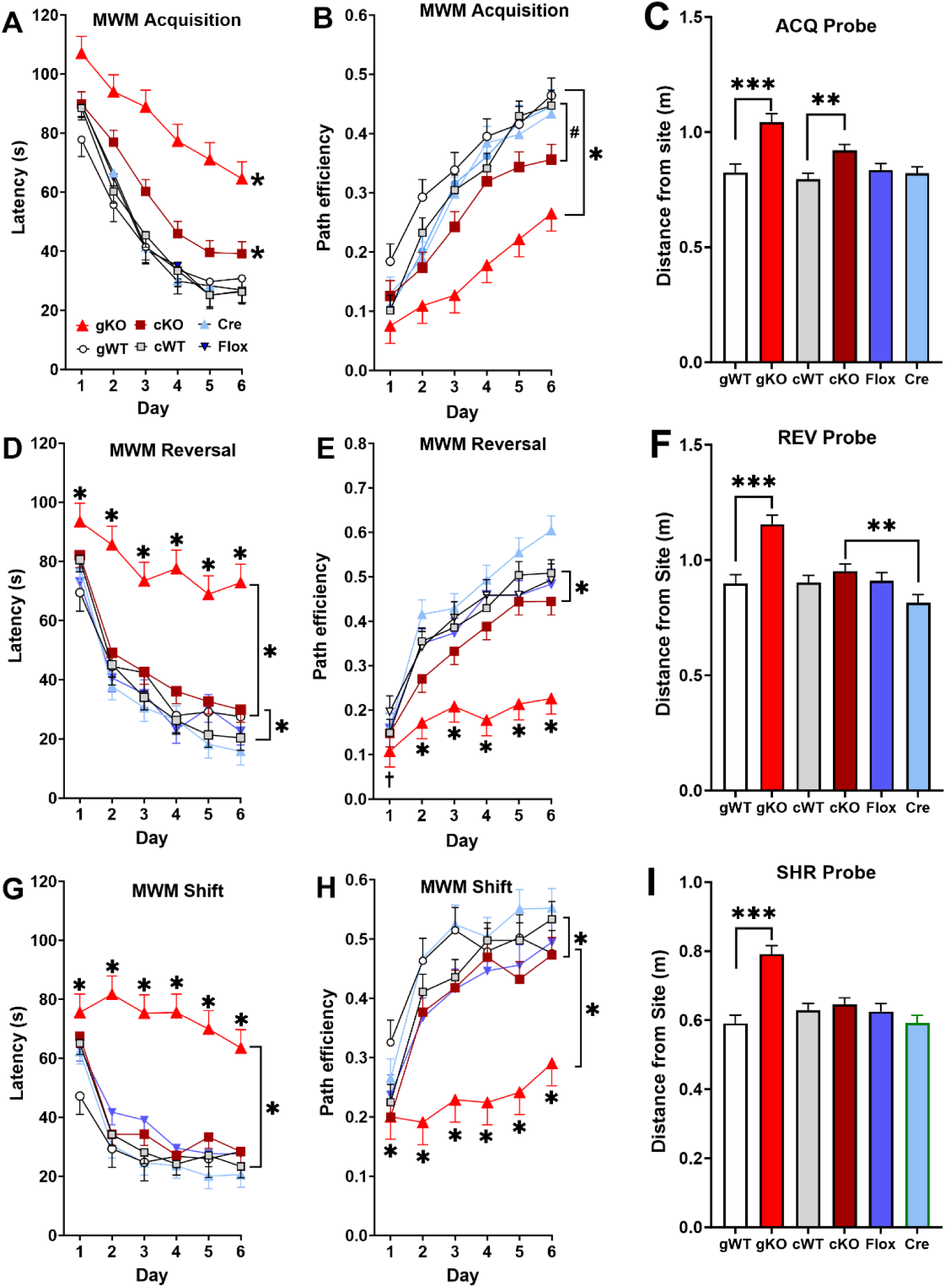
Morris Water Maze. A, Acquisition latency. **B,** Acquisition path efficiency. **C,** Acquisition probe average distance from the platform site. **D,** Reversal latency. **E,** Reversal path efficiency. **F,** Reversal probe average distance from the platform site. **G,** Shift latency. **H**, Shift path efficiency. **I**, Shift probe average distance to the platform site. Group sizes: cKO: 17 F, 16 M; cWT: 17 F, 14 M; Cre: 15 F, 13 M; Floxed: 14 F, 13 M; gWT: 15 F, 14 M, gKO: 15 F, 14 M.*P<0.05 **P<0.01, ***P<0.001.

On acquisition probe trials, gKO rats had increased average distance to the former platform site compared with gWT rats [F(1, 30.5) = 20.51, p<0.0001] (Fig. 9C). For target zone entries, there was a trend where gKO rats had fewer entries than gWT rats [F(1,30.7) = 3.14, p = 0.08]. The cKO rats had increased average distance to the former platform site compared with the other genotypes [Genotype: F(3,100)= 4,65, p<0.01] (Fig. 9C). There were no interactions with genotype.

For reversal latency, there was a main effect of genotype for gKO rats [F(1, 44.1) = 34.75, p<0.0001] (Fig. 9D) in which gKO rats had longer latencies than gWT rats. There was a genotype x day interaction [F(5, 245) = 3.85, p<0.01]. Slice-effect analyses showed longer latencies in gKO rats compared with gWT rats on all days, but the size of the difference varied, becoming larger across days. The gKO rats had decreased path efficiency compared with gWT rats [Genotype: F(1, 45.3) = 40.78, p<0.0001] (Fig. 9E). There was an interaction of genotype x day [F(5, 240)= 2.24, p<0.05] in which the magnitude of the deficit in gKO rats increased across days. There were no effects on swim speed; mean ± SEM were: gKO, 24.19 ± 0.69 cm/s; gWT, 23.46 ± 0.69 cm/s.

For cKO rats, there was a main effect of genotype [F(3, 151) = 2.71, p<0.05] (Fig. 9D). The cKO rats had longer latencies than cre rats, but there were no differences between cKO, flox, and cWT rats. There were no interactions with genotype. For cKO path efficiency, there was a main effect of genotype [F(3, 117) = 5.41, p<0.01] (Fig. 9E). The cKO rats had less efficient paths to the goal than cWT, cre, or flox rats; there were no interactions with genotype. There were no genotype effects on swim speed; mean ± SEM were: cKO, 23.89 ± 0.44 cm/s; cWT, 23.38 ± 0.44 cm/s; flox, 25.41 ± 0.49 cm/s; cre, 23.62 ± 0.48 cm/s.

For reversal probe trials, gKO rats had increased average distance to the former platform site compared with gWT rats [F(1, 48.7)= 34.27, p<0.0001] (Fig. 9F). For target zone entries, gKO rats had fewer entries compared with gWT rats [F(1, 51.1) = 11.05, p<0.01]. The cKO rats had increased average distance to the former platform site compared with cre rats but not compared with flox or cWT rats [Genotype: F(3, 85.4) = 3.79, p<0.05] (Fig. 9F). For target zone entries, there were no genotype effects.

For shift, gKO rats had longer latencies compared with gWT rats [genotype: F(1, 58.5) = 36.72, p<0.0001] (Fig. 9G). There was a genotype x day interaction [F(5, 250)= 3.68, p<0.01]. In gKO rats the longer latencies were significant on all days but became larger across days. The gKO rats had decreased path efficiency compared with gWT rats [F(1, 63.1) = 34.76, p<0.0001] (Fig. 9H). There was an interaction of genotype x day on path efficiency [F(5, 249) = 2.92, p<0.05]. The gKO rat differences varied as a function of day but were significant on all days and become larger across days. There were no genotype effects on swim speed; mean ± SEM were: gKO, 23.65 ± 0.67 cm/s; gWT, 22.73 ± 0.67 cm/s. For cKO rats, there was a main effect of genotype for latency [F(3, 171) = 2.59, p= 0.05] (Fig. 9H). Turkey’s post hoc tests showed a trend (p = 0.08) for floxed rats to have shorter latencies compared with cKO rats on days 2, 4, and 5.

No differences were seen between the other genotypes. For cKO rats, there was a main effect of genotype for path efficiency [F(3, 186) = 4.35, p<0.01] (Fig. 9H). Cre rats had greater path efficiency compared with floxed (p<0.05) or cKO rats (p<0.01) but did not differ from cWT rats. There were no genotype-related interactions. There was a main effect of genotype on swim speed [F(3,134)= 3.99,p<0.01]. Post hoc analyses showed an increase in swim speed for floxed rats compared with cre and cWT rats, but no differences compared with cKO rats; mean ± SEM were: cKO, 23.58 ± 0.44 cm/s; cWT, 23.19 ± 0.44 cm/s; flox, 25.07 ± 0.49 cm/s; cre, 23.16 ± 0.49 cm/s.

For shift probe trials, gKO rats had longer paths to the former platform site compared with gWT rats [F(1, 48.7) = 34.27, p<0.0001] (Fig. 9I). There was an interaction of genotype x day [F(1, 46.2) = 6.43, p<0.05] (not shown). Slice-effect analyses showed increased distance to the former platform site of gKO rats on both probe trials compared with gWT rats, but the effect was larger on the second probe trial. For target site entries, gKO rats had fewer entries compared with gWT controls [Genotype: F(1, 25.9)= 12.95, p<0.001]. There were no genotype interactions. For cKO rats there was no main effect of genotype for average distance to the target site (Fig. 9I). There were no genotype-related interactions. For target site entries, there were no genotype main effects or interactions.

For cued-random trials with curtains closed around the tank, gKO rats had longer latencies to the visible platform compared with gWT rats [F(1, 35.8) = 10.84, p<0.01] (Fig. 7D). There was no genotype x trial interaction. For cKO rats, there were no effects of genotype or genotype x trial interactions.

### CWM-Mirror

For latency, gKO rats took longer to escape compared with gWT rats [F(1, 40.4) = 25.05, p<0.0001] (Fig. 10A,B). There were no interactions with genotype. For cKO rats, there was a main effect of genotype [F(3, 107) = 9.62, p<0.0001] (Fig. 10A,B). cKO, cre, and floxed rats had longer latencies compared with cWT rats.

**Figure 10.**
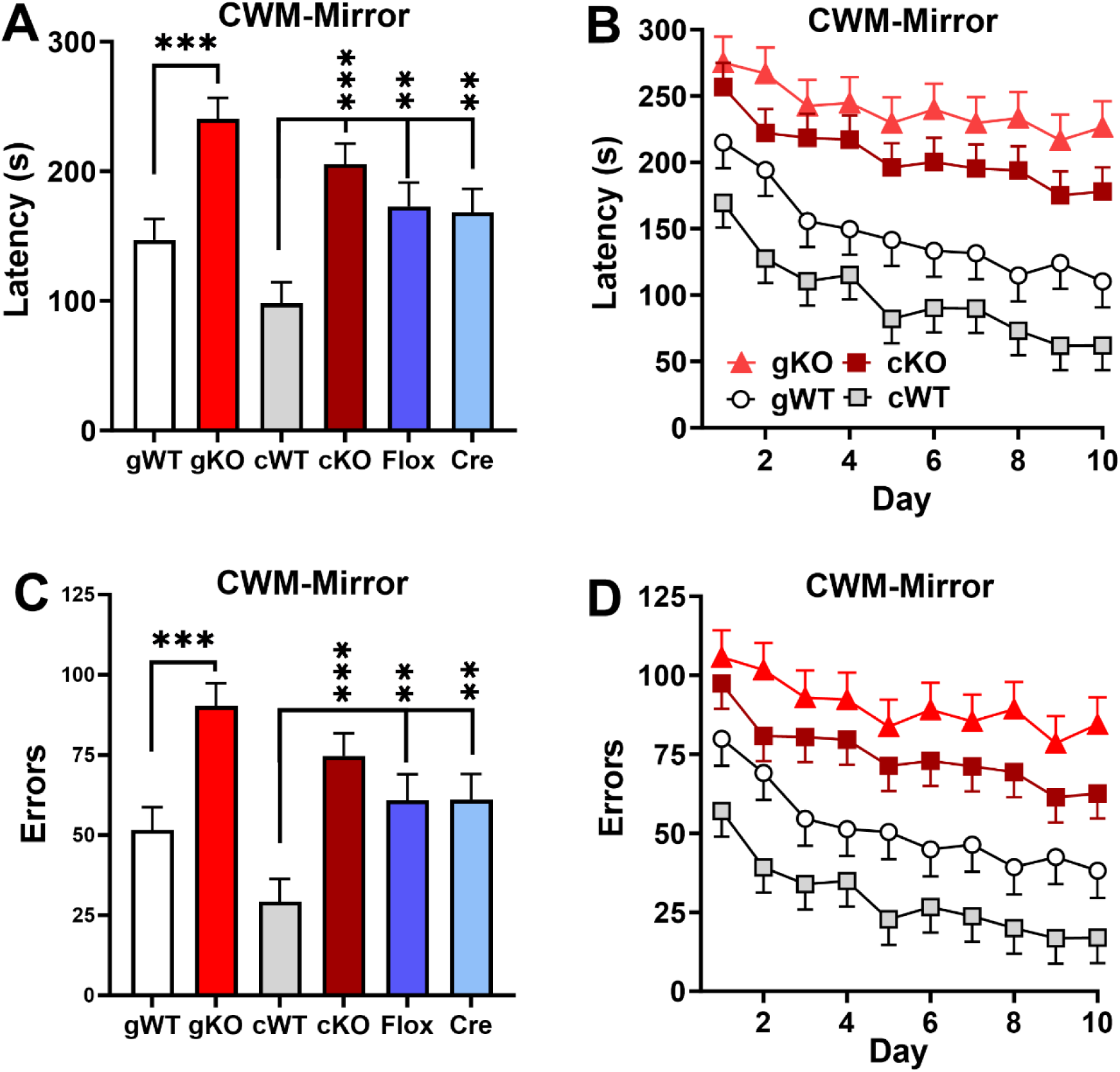
**CWM-mirror**. **A,** Latencies averaged across days. **B,** Latencies by day. **C,** Errors averaged across days. **D,** Errors by day. Group sizes: cKO: 17 F, 16 M; cWT: 17 F, 14 M; Cre: 15 F, 13 M; Floxed: 14 F, 13 M; gWT: 15 F, 14 M, gKO: 15 F, 14 M. **P<0.01, ***P<001.

The gKO rats made more errors compared with gWT rats [Genotype: F(1, 1.7) = 23.56, p<0.0001] (Fig. 10C,D). There was an interaction of genotype x day [F(9, 452) = 1.90, p<0.05]; the gKO group had increased errors on all test days and the size of the difference increased across days. For cKO there was also a genotype effect [Genotype: F(3, 105) = 8.82, p < 0.0001] (Fig. 10C,D). cKO, cre, and floxed rats had increased errors compared with cWT rats (cre and flox groups not shown for clarity). There were no genotype-related interactions.

### qRT-PCR

To estimate the efficacy of the cre-lox system to delete *Lphn3* in TH-expressing cells, qPCR was used to show a decrease in *Lphn3*. There was a main effect of genotype [F(3, 47.7) = 5.94, p<0.001] where *Lphn3* cKO rats had decreased expression compared with cWT rats (Fig. 11). There was no effect of sex or interaction of genotype x sex. Immunohistochemistry showed a colocalization of TH and cre-dependent tdTomato virus, showed that the cre targets TH expressing cells (Fig. 12)

**Figure 11.**
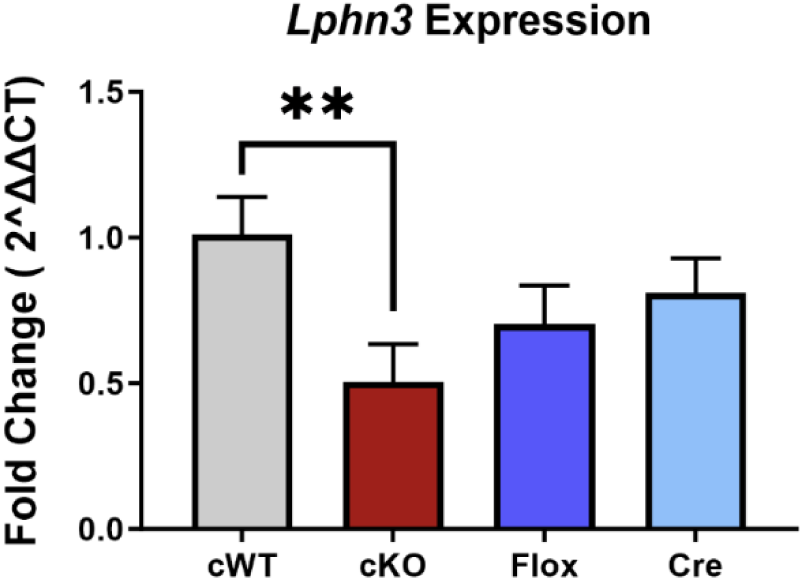
Real Time Polymerase Chain Reaction (RT-PCR): *Lphn3* cKO rats had a reduction in *Lphn3* in the caudate putamen. Group sizes: cWT: 8 F, 8 M; Flox: 8 F, 8 M; Cre: 8 F, 8 M; cKO: 8 F, 8 M. **P<0.01.

**Figure 12:**
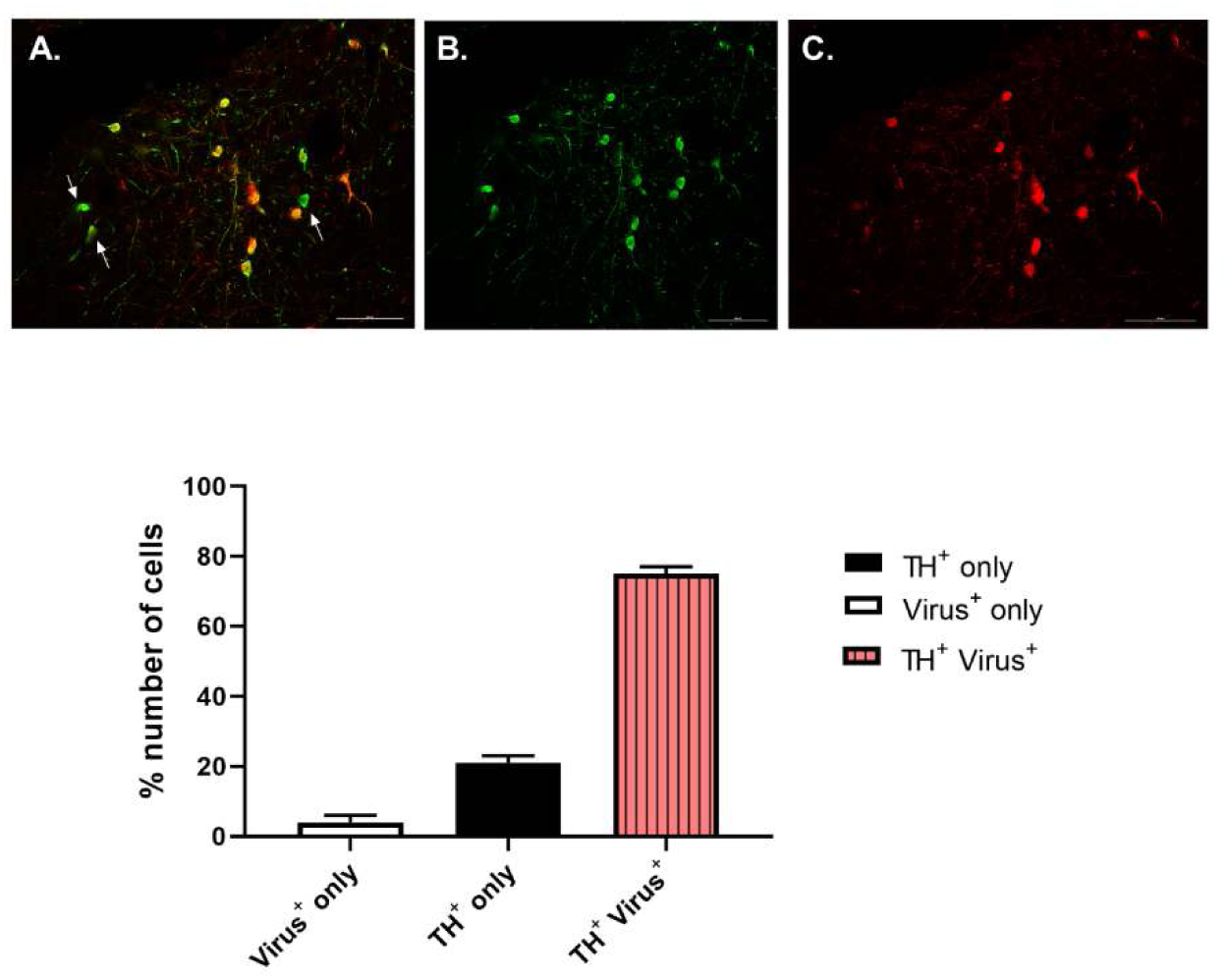
Fluorescent immunohistochemistry analysis of TH and TdTomato in the striatum. A, Merged image. **B,** DAPI. **C,** TH. **D,** TdTomato virus. DAPI = blue. TH = Green. Virus = Red. Scale = 100 µm. Lower panel: quantification in 100 cells.

## DISCUSSION

We previously created a constitutive/global *Lphn3* KO (gKO) and showed it has a pronounced phenotype (Regan, 2019; Regan et al., 2020; Regan et al., 2021b; Regan et al., 2021a; Regan et al., 2022). *Lphn3* gKO rats have dysregulated striatal dopamine release (Regan et al., 2020), increased levels of striatal TH, L-aromatic amino acid decarboxylase, and DARPP-32 with reduced levels of the DA D1 receptor and transporter (DAT) by western blot in combination with hyperactivity (Regan et al., 2019). The gKO rats also exhibit large deficits in egocentric and allocentric learning and memory (Regan et al., 2021b) and deficits in delayed spatial alternation and differential reinforcement of low rate responding (Sable et al., 2021) but no differences in impulsive choice (Carbajal et al., 2023).

LPHN3 is a complex protein. Its extracellular domain binds to presynaptic adhesion molecules to form trans-synaptic bridges. Its highest affinity is for FLRT3, but it can also bind with Tenurin2 (Wang et al., 2024). These bridges are thought to modulate synaptic activity. LPHN3 has an autocleavage site that forms alternate splicing types, either Gα_s_ or Gα_12/13_, wherein Gα_s_ facilitates synapse formation (Wang et al., 2024). There are between 15 (Acosta et al., 2016) and 21 variants (Domene et al., 2011b) of *LPHN3* associated with ADHD but which of these are the most important or how they alter the synaptic function of LPHN3 containing neurons is unknown.

The behavioral deficits of *Lphn3* gKO rats can be partially explained by LPHN3 expression. LPHN3 is expressed in multiple brain regions, with highest expression in the striatum and nucleus accumbens with lower expression in cerebellum and hippocampus. In hippocampus, it is expressed primarily in CA1 and dentate gyrus with some expression in the PFC. In CA1, LPHN3 occurs in excitatory neurons (Sando et al., 2019a) and this may account for the larger effect of the gKO on MWM performance compared with that of the cKO. Sando et al., showed that the loss of *Lphn3* creates a deficit in excitatory glutamatergic neuronal function. In the cerebellum, Zhang et al., found that loss of only one, *Lphn3* or *Lphn2*, had no effect on EPSCs in Purkinje cells whereas simultaneous loss of both reduced EPSC currents by 50%; loss in other cerebellar neurons has no effect (Zhang and Liakath-Ali, 2020). This project determined if the deletion of *Lphn3* selectively in TH positive cells confers a phenotype different from that of the gKO rats. The results showed differences and similarities between the gKO and cKO rats.

The gKO rats were hyperactive compared with gWT whereas cKO rats were not in a 72-h test of locomotor activity. Rodent activity generally peaks 5-6 h into the night phase (Zoratto et al., 2013) and this peak was heightened in gKO rats. Once hyperactivity emerged, it reemerged during each dark cycle and was absent during light cycles.

Egocentric/procedural learning and memory depends on striatal DA, as demonstrated using 6-OHDA induced striatal lesions (Braun et al., 2015; Braun et al., 2016; Vorhees and Williams, 2016). The gKO rats have large deficits in egocentric learning in the CWM (Regan et al., 2021a). The cKO rats have a similar deficit in the CWM. However, on the reverse, mirror image CWM test, both the gKO and cKO rats had impaired learning with the cKO deficit not as large as that seen in the gKO rats.

Allocentric spatial learning and memory is dependent on hippocampus and entorhinal cortex (Buzsaki and Moser, 2013; Ekstrom et al., 2014; Buzsaki and Llinas, 2017). The gKO rats showed striking deficits in spatial learning in the MWM on acquisition and on reversal and shift phases that assess cognitive flexibility. They also showed impaired reference memory on probe trials a day after each learning phase. By contrast, cKO rats showed only small deficits in the MWM across phases, and a probe trial reference memory deficit only at the end of acquisition but not after reversal or shift trials. If LPHN3 is predominately expressed in the striatum, the larger effect of the cKO on egocentric navigation in the CWM compared with the smaller effect on allocentric navigation in the MWM would be expected. Tempering this interpretation are data showing that LPHN3 is located in the CA1 region of the hippocampus where it modulates activity in glutamatergic neurons by acting on NMDA receptors (Sando et al., 2019b). Perhaps the role of LPHN3 in the hippocampus is less than in the striatum or not expressed in hippocampal TH neurons but still affects spatial learning and memory in the MWM albeit to a lesser degree.

The RWM is also thought to rely on DA (Bernhardt et al., 2018). The *Lphn3* cKO rats, however, had no differences in RWM performance compared with cWT rats. The lack of effect of *Lphn3* deletion in both mouse and rat models could be because the water version of the RWM is, perhaps, less sensitive of a test compared with the CWM and MWM tests. *Lphn3* is also expressed in the PFC but at lower levels than in striatum and other regions. *Lphn3* in the PFC may function in concert with *Lphn1 or 2* as occurs in the cerebellum such that deleting LPHN3 in the PFC is insufficient to elicit as clear phenotype. Interestingly, *Lphn3* KO mice performed similarly to WT mice in a working memory task (Orsini et al., 2016); since working memory depends on the PFC these data suggest that LPHN3 has little or no role in working memory.

A limitation of our study is that deletion of *Lphn3* is not complete in TH neurons. Although *Lphn3* is decreased within the striatum (Fig. 11), there is co-expression of the viral cre reporter and TH antibody (Fig. 12), cre-recombinase does not capture 100% of *Th*-expressing cells. Nevertheless, it is notable that even the partial deletion of *Lphn3* in the cKO rats produced a phenotype consistent with that of the gKO rat. Based on the results, we predict that complete removal of *Lphn3* in TH-expressing cells would produce an even more pronounced phenotype.

The data show that LPHN3 plays a role in DA related behaviors and by comparison of the gKO and cKO rats, it can be inferred that LPHN3 also has a role in glutamatergic signaling in the hippocampus.

